# The multidisciplinary nature of COVID-19 research

**DOI:** 10.1101/2020.11.23.394312

**Authors:** Ricardo Arencibia-Jorge, Lourdes García-García, Ernesto Galbán-Rodríguez, Humberto Carrillo-Calvet

## Abstract

**Objective:** We analyzed the scientific output after COVID-19 and contrasted it with studies published in the aftermath of seven epidemics/pandemics: Severe Acute Respiratory Syndrome (SARS), Influenza A virus H5N1 and Influenza A virus H1N1 human infections, Middle East Respiratory Syndrome (MERS), Ebola virus disease, Zika virus disease, and Dengue.

**Design/Methodology/Approach:** We examined bibliometric measures for COVID-19 and the rest of studied epidemics/pandemics. Data were extracted from Web of Science, using its journal classification scheme as a proxy to quantify the multidisciplinary coverage of scientific output. We proposed a novel Thematic Dispersion Index (TDI) for the analysis of pandemic early stages.

**Results/Discussion:** The literature on the seven epidemics/pandemics before COVID-19 has shown explosive growth of the scientific production and continuous impact during the first three years following each emergence or re-emergence of the specific infectious disease. A subsequent decline was observed with the progressive control of each health emergency. We observed an unprecedented growth in COVID-19 scientific production. TDI measured for COVID-19 (29,4) in just six months, was higher than TDI of the rest (7,5 to 21) during the first three years after epidemic initiation.

**Conclusions:** COVID-19 literature showed the broadest subject coverage, which is clearly a consecuence of its social, economic, and political impact. The proposed indicator (TDI), allowed the study of multidisciplinarity, differentiating the thematic complexity of COVID-19 from the previous seven epidemics/pandemics.

**Originality/Value:** The multidisciplinary nature and thematic complexity of COVID-19 research were successfully analyzed through a scientometric perspective.

## Introduction

Epidemics and pandemics have significant repercussions in human societies, spread beyond national borders, and affect many people in extensive areas (Porta Serra, 2014). Throughout history, humankind has suffered millions of deaths due to these diseases worldwide, undergoing vast economic and social losses (Fan, Jamison, and Summers, 2018; Huber, Finelli, and Stevens, 2018; Keogh-Brown and Smith, 2008; Kuhar and Fatović-Ferenčić, 2020). The early implementation of suppression or mitigation strategies has been crucial to face adverse effects (Kuhar and Fatović-Ferenčić, 2020; Pike et al., 2014). In such contexts, the integration of various sectors of society in decision-making processes has been paramount to balance health and economic priorities and minimize the impact in almost every aspect of social life.

COVID-19 pandemic is a disruptive experience for everyone, including scientists (Myers et al., 2020). The massive increase in scientific articles might have overcome researchers. Dozens of bibliometric studies have been published after the emergence and growing expansion of the new pandemic. Most of them include traditional bibliometric indicators, such as main contributing authors, journals, institutions, and countries (Chahrour et al., 2020; Darsono, Rohmana, and Busro, 2020; De Felice and Polimeni, 2020; Dehghanbanadaki et al., 2020; Kambhampati and Vaish, 2020). Thematic clusters have been identified through mapping techniques according to keywords co-occurrence, co-citation patterns, or international collaborations (El Mohadab, Bouikhalene, and Safi, 2020; Hamidah, Sriyono and Hudha, 2020; Herrera-Viedma et al., 2020). A few of them have compared bibliometric patterns of epidemic/pandemic diseases (Tao et al., 2020; Zhai et al., 2020; Zhang et al., 2020; Zhou and Chen, 2020). However, we consider that previous reports have not been focused on research multidisciplinarity.

The current study aims to analyze and compare scientific literature multidisciplinarity on COVID-19 and other 21st century epidemics/pandemics using a novel Thematic Dispersion Index (TDI). We consider that we successfully overcame the limitations of previous reports.

## Material and methods

### Data Recovery

Scientific output data of each epidemic/pandemic was retrieved from the Web of Science™ (WoS, developed by Clarivate Analytics) on June 30, 2020, with the following search strategies: COVID: TS=COVID OR TS=SARS-COV-2; EBOLA: TS=EBOLA; DENGUE: TS=DENGUE; H1N1: TS=(“Influenza A”) AND TS=H1N1; H5N1: TS=(“Influenza A”) AND TS=H5N1; MERS: TS=(“Middle East Respiratory Syndrome”) OR TS=(“MERS-COV”); SARS: TS=(“sars-cov”) NOT TS=(“sars-cov-2”) OR TS=(“Severe Acute Respiratory Syndrome”) NOT TS=(“Severe Acute Respiratory Syndrome Coronavirus 2”); ZIKA: TS=ZIKA.

Search dates ranged as follows for the different diseases: 2003 to 2005 for Severe Acute Respiratory Syndrome (SARS); 2005 to 2007 for Influenza A virus H5N1 human infection; 2009 to 2011 for Influenza A virus H1N1 human infection; 2012 to 2014 for Middle East Respiratory Syndrome (MERS); 2014 to 2016 for Ebola and Zika virus diseases; 2015 to 2017 for Dengue; and January to June 2020 for COVID-19.

### Procedure

Multidisciplinarity was addressed from a bibliometric perspective, using the WoS journal classification scheme as a proxy (Leydesdorff and Bornmann, 2016). According to the WoS classification scheme, a subject category is assigned to each journal according to its subject scope. Hence, journal subject classification was selected due to its higher aggregation, with the journal being more multidisciplinary as the number of subject categories increase.

Likewise, citations received by this set of articles were taken into account, as a measure of impact on the scientific community (Garfield, 2006). The multidisciplinary nature will also be expressed if journals from citing articles belong to an increasing number of subject categories.

### Basic Indicators

The following indicators were calculated for the set of articles of each epidemic/pandemic:

**A:** articles published during the first three years (only the first six months were considered for COVID-19); **A Cit:** citing articles during the first three years after the outbreak; **Cit (mean):** mean of citations received in one year by papers published during the same year (calculated for the first, second and third year after disease emergence); **WCs:** Web of Science subject categories; **Cit WCs:** Citing Web of Science subject categories.

### New index proposed

Following the Pareto principle, a Thematic Dispersion Index (**TDI**) was developed. It aimed to balance the core of WoS subject categories (WCs) that comprise 80% of articles (Thematic Concentration of Scientific Production, **TCp**), and 80% of citing articles (Thematic concentration of citations, **TCc**) (Arencibia-Jorge, Vega-Almeida, and Carrillo-Calvet, 2020), according to the following formula:

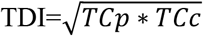

TDI has the following property: TDI ≥ 1 < 1x, where: x equals the maximum number of WCs (currently 255). Values close to 1 express a high degree of disciplinary specialization or scientific production concentration, while higher values will increasingly determine multidisciplinary nature.

## Results

Despite the international scientific community reaction given the dramatic explosion of literature (more than 8 000 papers covered by WoS during the first six months, Figure 1a and 1c), the dynamic patterns of COVID-19 research could be qualitatively similar to those observed after other 21st century epidemics/pandemics. We have confirmed that emerging infectious diseases with an epidemic/pandemic nature typically unleash an accelerated growth of scientific production during the three years after outbreak emergence (Figure 1b). This initial stage generates an avalanche of data and evidence derived from the damaging effects of the disease and the growth of incentives to face the problem. Subsequently, once the health emergency is under control or effective treatments have appeared, a gradual decrease in scientific production begins.

**Figure 1.**
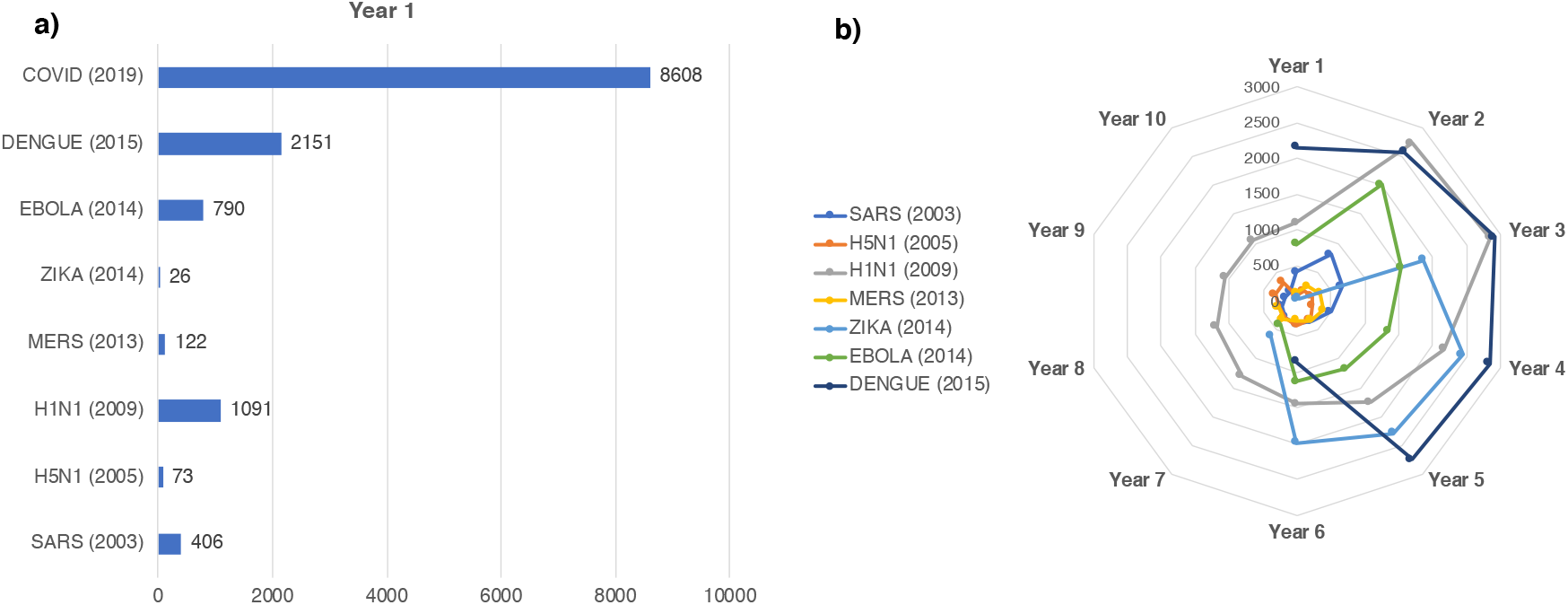
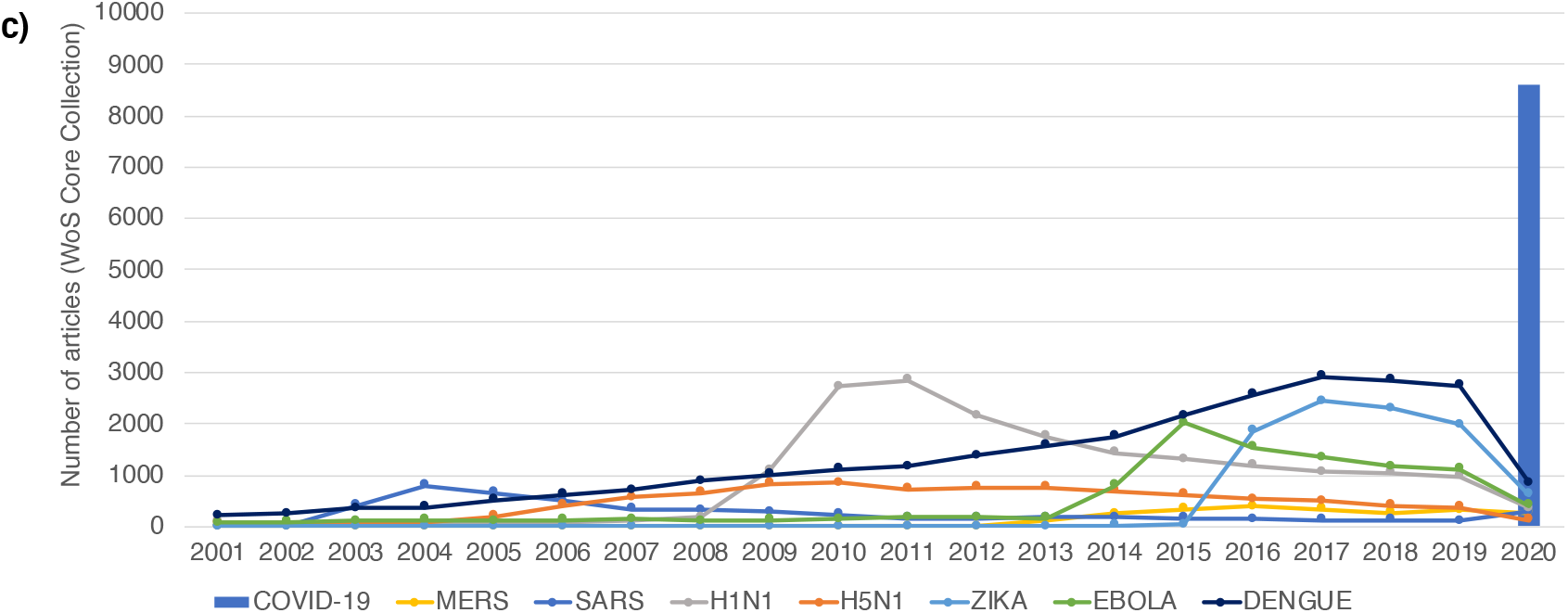
Evolution of the scientific production on eight pandemic diseases of the 21st century. **a)** first year of epidemic/pandemic emergence; **b)** ten years after epidemic/pandemic emergence; **c)** during 2001-2020. (Source: WoS. Date of retrieval: June 30, 2020).

Likewise, the average annual citations received by articles published in those first three years will increase, reaching its maximum peak generally the third year after outbreak emergence (Table 1). This phenomenon has occurred regularly during each of the seven epidemics/pandemics that preceded COVID-19. However, one remarkable aspect of the new pandemic is the multidisciplinary nature of research.

**Table 1.** Bibliometric measures of productivity, impact, and multidisciplinary scope after selected epidemics/pandemics (Source: WoS. Date of retrieval: June 30, 2020).

| Epidemics/<br>Pandemics<br>(Start Year) | A | A Cit | Cit<br>(mean)<br>Year 1 | Cit<br>(mean)<br>Year 2 | Cit<br>(mean)<br>Year 3 | WCs | Cit<br>WCs | TCp | TCc | TDI |
| --- | --- | --- | --- | --- | --- | --- | --- | --- | --- | --- |
| SARS (2003) | 1871 | 3272 | 4.08 | 9.25 | 15.21 | 144 | 163 | 18 | 18 | 18.0 |
| H5N1 (2005) | 453 | 1858 | 3.41 | 9.01 | 14.00 | 71 | 139 | 9 | 11 | 9.9 |
| H1N1 (2009) | 3694 | 7628 | 2.69 | 6.27 | 10.9 | 134 | 173 | 11 | 13 | 12.0 |
| MERS (2012) | 712 | 1368 | 2.92 | 9.73 | 12.57 | 69 | 93 | 7 | 8 | 7.5 |
| EBOLA (2014) | 4344 | 6661 | 1.48 | 3.61 | 7.64 | 176 | 197 | 20 | 22 | 21.0 |
| ZIKA (2014) | 1934 | 1930 | 1.04 | 4.41 | 6.50 | 130 | 130 | 13 | 13 | 13.0 |
| DENGUE (2015)* | 7636 | 13748 | 0.69 | 4.62 | 8.07 | 169 | 205 | 15 | 22 | 18.2 |
| COVID-19 (2020) | 8608 | 8501 | 3.53 | - | - | 203 | 188 | 32 | 27 | 29.4 |

\*Dengue has had pandemic characteristics during this century. More than 4000 deaths were reported in 2015. The largest number of dengue cases ever reported globally was in 2019. All regions were affected.

Five of the seven severe outbreaks before COVID-19 involved journal articles covering more than one hundred Web of Science subject categories (WCs). In the five cases, the thematic core of scientific production encompassed 10 to 25 WCs, mainly connected to biomedical domains (Table 1; Figure 2).

**Figure 2.**
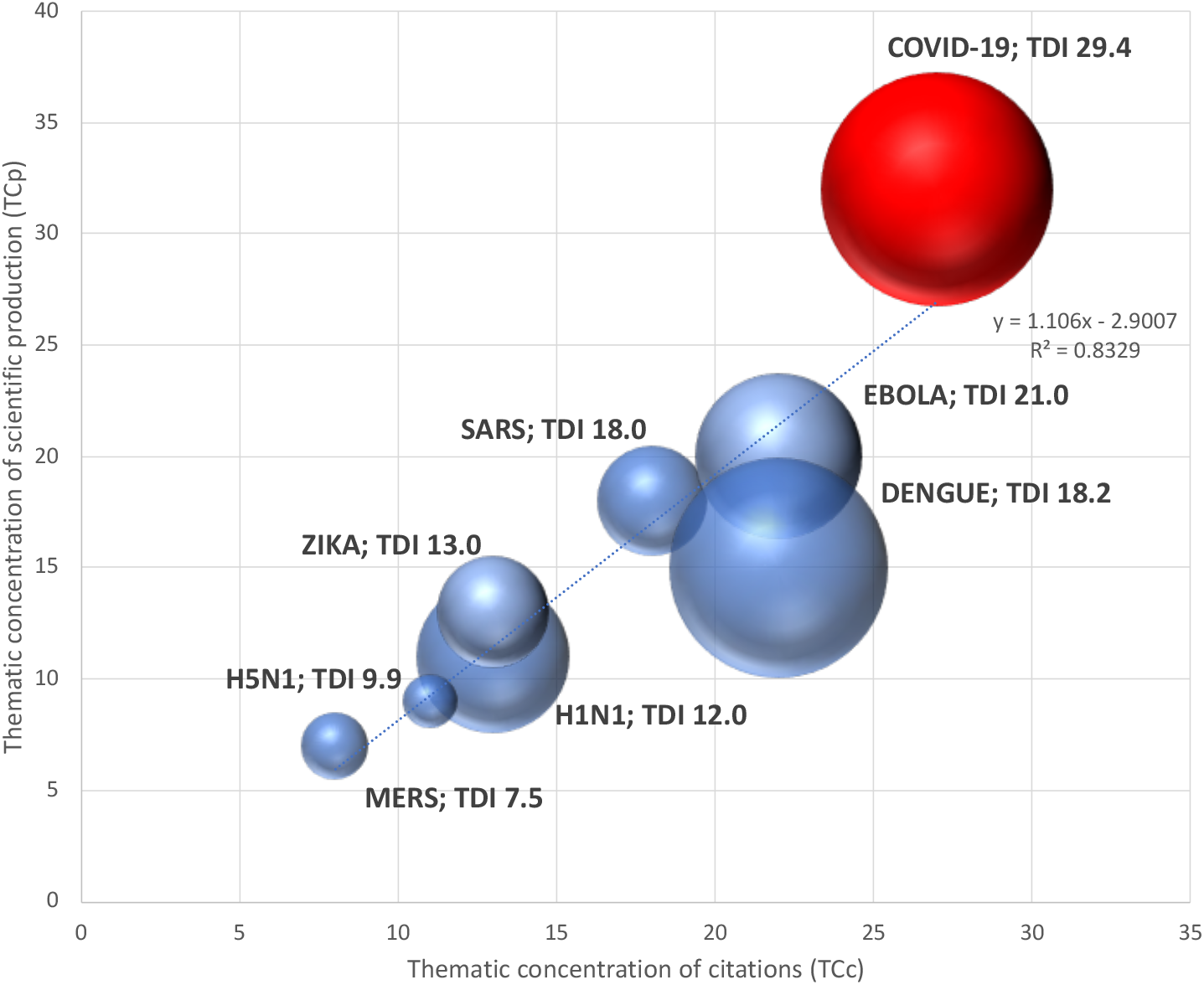
Thematic concentration of articles and citations, and thematic dispersion index (TDI) of the eight analyzed pandemics during the first three years after disease emergence. Bubble’s size is proportionate to TDI. Blue bubbles: Severe Acute Respiratory Syndrome (SARS); Influenza A virus H5N1 and Influenza A virus H1N1 human infections, Middle East Respiratory Syndrome (MERS), Ebola virus disease, Zika virus disease, and Dengue. Red bubble: COVID-19.

Therefore, research related to epidemic/pandemic diseases tends to go beyond disciplinary frameworks. We observed that after the eight epidemics/pandemics, the multidisciplinarity was related to the disease mortality, biomedical complexity, global dispersion, and economic impact. Hence, Influenza A-H5N1 and MERS, which affected fewer countries, had a lower multidisciplinary scope and concentrated the least number of articles published during the three years after outbreak emergence.

COVID-19, in a short time, has become the most extensive and deadliest pandemic of this century (more than 500,000 deaths during the first semester, and two million expected for the next year). COVID-19 research has increasingly involved different thematic domains (more than two hundred WCs) directly or indirectly related to the disease, prevention, diagnosis, treatment, and social response. The TDI reached by the current pandemic is the highest of all analyzed. It has nearly 30 WCs concentrating on the most massive volumes of articles and citations, reflecting the multidimensional impact that the disease has had in just six months.

## Discussion

Bibliometric multidisciplinary, interdisciplinary, or transdisciplinary analyses use two main approaches: A Bottom-up approach, based on clustering sets of articles according to a bibliographic criterion, either bibliographical coupling or co-citation networks; and a Top-to-bottom approach dependent on existing classifications schemes (Wagner et al., 2011). These strategies search for communication channels between authors, institutions, cited or citing authors, cited or citing journals, cited or citing documents, and mapping techniques to illustrate the diversity of research areas (Chen, 2017; Klavans and Boyack, 2011; Mochini et al., 2020).

In just a few months, bibliometric indicators of COVID-19 have surpassed by far what has been observed in other epidemics/pandemics. There are five WoS subject categories (Medicine General & Internal, Infectious Diseases, Virology, Immunology, and Microbiology) shared by the thematic core of eight studied pandemics. COVID-19 research hotspots have dealt with molecular virology, immunogenetics, epidemiology, medicine, imaging, pharmacology, environmental sciences, economics, anthropology, social sciences, philosophy, ethics, and other disciplines (El Mohadab, Bouikhalene, and Safi, 2020). Our bibliometric analysis demonstrated the growing thematic expansion with COVID-19.

Several reasons explain the multidisciplinary burst of COVID-19 research, as compared to other epidemics/pandemics. Its global distribution, incidence rates initially affecting high-income countries, high fatality rates in certain groups, wide-ranging clinical manifestations, lack of a vaccine or specific treatment, and psychological, social, political, and economic consequences, may altogether motivate the keen interest of a vast scientific community. Probably, the current technological resources plus extensive usage of social media and pre-print communications have also favored and accelerated communication between scientists from different disciplines and countries.

At the same time, COVID-19 is a respiratory infectious disease with numerous extrapulmonary manifestations that turn it into a multi-organ condition (Wang et al., 2020). Patients suffering from the most frequent chronic diseases have been significantly affected. Diabetes, hypertension, cardiovascular diseases, and cancer account for a highly vulnerable population with a worse prognosis contributing to death rates statistics. Hence, clinical consequences have been varied and severe, requiring the conjunction of different medical specialties. Multispecialty is a hallmark of COVID-19 research published in journals from various medical fields, seldom seen in previous epidemics/pandemics.

Additionally, this pandemic has strong links with the environment and OneHealth approaches. Leading approaches have included the origin of the virus, wildlife surveillance, risk reduction, and biosecurity. These topics have required the integration of researchers from Biological Sciences, such as Zoology, Genetics, Evolutionary Biology, Physiology, Biochemistry, and Cell Biology, in collaboration with ecologists, physicians, epidemiologists, and even in occupational health and safety specialists. Lockdown strategies have also required economists, sociologists, psychologists, psychiatrists, and researchers from Management, Commerce, Transport, and Tourism.

The techno-driven approach based on artificial intelligence and machine learning used to forecast outbreaks resurgence, SIR modeling, imaging processing, and diagnosis tools have also contributed. Internet of things (IoT) devices, big data, robotics, drone technologies, and multiple mobile applications have also been used to monitor mobility and track active cases and contacts.

Multidisciplinary research and development (R&D) approaches are crucial to effectively face a pandemic (Peters, Jandrić, and McLaren, 2020; Moradian et al., 2020). With the incorporation of specialists from various multinational teams, better conditions are created to develop interdisciplinary and transdisciplinary research on COVID-19 (Moradian et al., 2020). Thus, data analysts, computer specialists, robotics experts, engineers from various branches, sociologists, psychologists, philosophers, lawyers, information scientists, among others, have provided input to COVID-19 research.

The diversification of journal ecosystems and peer-review acceleration (Smart, 2020), together with the open exchange of experimental data following the Open Access and the more recent Open Science movements, could have also contributed to the explosion of COVID literature (Belli et al., 2020). The intensive and accelerated use of pre-print servers has been another fingerprint of the new pandemic (Kozlakidis et al., 2020; Johanson et al., 2020). Overall, we testify an unprecedented phenomenon that just started, and we are unable to predict its extent and magnitude. Nevertheless, we consider that the productivity, impact, and multidisciplinary scope of COVID-19 literature has been extraordinary and will continue to grow for at least three more years.

### Limitations

The main limitation of this study is its intrinsic bias. WoS-based results may differ from those obtained from other databases. However, previous studies have successfully used this data set (Zhang et al., 2020). To our knowledge, this is the first study that uses a journal classification scheme to analyze multidisciplinarity of COVID-19 from the bibliometric perspective. We consider that the WCs core offered an essential multidisciplinarity dimension as it covers the largest volume of articles and citations. The new metric we are proposing may be used to compare different levels of aggregation (institutions, individuals, countries), using complementary diversity measures to analyze the sets of publications (Moschini et al., 2020; Porter and Rafols, 2009). TDI would be relatively easy to implement in other internet-available databases with large journal classification schemes (e.g., Scopus or Dimensions).

## Conclusions

In only six months, COVID-19 literature involved increasing thematic domains directly or indirectly related to the disease and its social, economic, and political consequences. The newly proposed indicator (TDI), based on a database source-aggregated classification system, allowed the study of multidisciplinarity, differentiating the thematic complexity of COVID-19 from the previous seven epidemics/pandemics.

## Acknowledgements

This research was supported by the project “Scientometrics, Complexity and Science of Science”, at the Complexity Science Center of the National Autonomous University of Mexico (UNAM).

